## Supplemental Tables 1-6 for "Heat Shock Factor 1 forms nuclear condensates and restructures the yeast genome before activating target genes"

**Supplemental File 1**

Heat Shock Factor 1 forms nuclear condensates

and restructures the yeast genome

before activating target genes

Linda S. Rubio, Suman Mohajan and David S. Gross

Supplemental Tables

### Table 1. Yeast Strains

| **Strain name** | **Genotype** | **Source** |
| --- | --- | --- |
| BY4741 | *MATa his3Δ1 leu2Δ0 met15Δ0 ura3Δ0* | Research Genetics |
| W303-1B | *MATα ade2-1 trp1-1 can1-100 leu2-3,112 his3-11,15 ura3-1* | R. Rothstein |
| ASK727 | *MATa/MATα ura3-1/ura3-1 ade2-1/ade2-1 leu2-3,112/leu2-3,112 can1-100/CAN1 THR/thr1-4 trp1-1/trp1^-^ his3-11,15::GFP-LacI::HIS3/ his^-^ HSP104- LacO_256_::TRP1/HSP104 TMA10-TetO_200_::LEU2/TMA10 leu2::TetR-mCherry::hphMX::leu2 SEC63-Myc×13/SEC63* | (Chowdhary et al., 2019) |
| ASK741 | *MATa/MATα his3∆1/his3∆1 leu2∆0/leu2∆0 met15∆0/met15∆0 ura3∆0/ura3∆0 HSF1-GFP:: HIS3MX6/HSF1-GFP::HIS3MX6* | (Chowdhary et al., 2022) |
| DBY1447 | *MATa/MATα LacI-GFP:LEU2/leu2-3 trp1-1/trp1-1 can1-100/can1-100 ade2-1/ade2-1 p6LacO-SSA2/SSA2 p6LacO-SSA4/SSA4  LacI-GFP:HIS3/his3-1  SEC63-13myc:KANMX/SEC63  HTA2-mCherry:HIS5/HTA2* | Donna and Jason Brickner |
| DPY1561 | *MATa trp1-1 can1-100 leu2-3,112 his3-11,15 ura3-1 ADE2 HSF1-mVenus::HIS3 SIS1-mKate::URA3 HSP104-mTagBFP2::LEU2* | (Feder et al., 2021) |
| LRY031 | W303-1B x DPY1561 | This Study |
| LRY032 | *MATa/MATα A*de+ *trp1-1/ trp1-1 can1-100/can1-100 leu2-3,112/leu2-3,112 his3-11,15/his3-11,15 ura3-1/ura3-1 SIS1/SIS1-mKate::URA3 HSP104/HSP104-mTagBFP2::LEU2* | This Study |
| LRY033 | LRY032*; HSF1/HSF1-mNeonGreen::HIS3* | This Study |
| LRY037 | W303-1B; *HSF1-mNeonGreen::HIS3* | This Study |
| SCY004 | *MATa ADE2 trp1-1 can1-100 leu2-3,112 his3-11,15 ura3-1 HSF1-mVenus::HIS3 RPB3-mCherry::hphMX6* | (Chowdhary et al., 2022) |
| LRY040 | LRY037; *RPB3-mCherry::hphMX6* | This Study |
| LRY016 | *MATα ade2-1 trp1-1 can1-100 leu2-3,112 his3-11,15 ura3-1 LEU2::pGPD1-osTIR1* | (Rubio and Gross, 2023) |
| LRY100 | LRY016, *HSF1-mAID-9cMyc::HYG-MX* | This Study |
| LRY102 | LRY016, *RPB1-mAID-9cMyc::HYG-MX* | This Study |
| VPY705 | *MATa/MATα ade2-1/ade2-1 can1-100/can1-100 trp1-1/trp1-1 leu2-3,112/leu2-3,112 his3-11,15::GFP-LacI::HIS3/his3-11,15::GFP-LacI::HIS3 ura3-1/ura3-1::MCP-mCherry::URA3 HSP12-LacO_128_::ura3FOA^r^/HSP12 pHSP104-MS2×24::HSP104-LacO_256_::TRP1/HSP104 SEC63-Myc×13::TRP1/SEC63-MYC×13* | This Study |

### Table 2. Plasmids

| **Plasmid Name** | **Feature** | **Source** |
| --- | --- | --- |
| pFA6a-link-ymNeonGreen-SpHis5 | link-mNeonGreen-SpHIS5 | (Botman, de Groot, Schmidt, Goedhart, & Teusink, 2019) |
| pGZ154 | pGPD-OsTIR1-LEU2 | C.K. Govind |
| pHyg-AID*-9myc | miniAID-9MYC-HYGR | (Morawska & Ulrich, 2013) |

### Table 3. Primers used for Strain Construction

| **Name** | **Sequence (5’ 🡪 3’)** | **Purpose** |
| --- | --- | --- |
| Hsf1 ymNeonGreen F | AGGACCCGACAGAGTACAACGATCACCGCCTGCCCAAACGAGCTAAGAAAGGTGACGGTGCTGGTTTA | Tagging of Hsf1 with mNeonGreen (mNG) |
| Hsf1 ymNeonGreen R | ATACTATATTAAATGATTATATACGCTATTTAATGACCTTGCCCTGTGTATCGATGAATTCGAGCTCG |  |
| HSF1_Conf 2 _F | ACGACAATAACACTAGTGAGG | Confirmation of Hsf1-mNG tagging |
| HSF1_Conf 2_R | CTCAGGCTCTCACTAGCTC |  |
| RPB3 C-term F | TATTGTTAGCTCTGACACAGATG | Amplification of *RPB3-mCherry* |
| RPB3 C-term R | GTAGATTTGACATTCGGTAGTTC |  |
| RPB3 Conf F | GATCAAGTTGTCGTCAGAGGTATCG | Confirmation of Rpb3-mCherry tagging |
| RPB3 Conf R | ATTAGTAGACGAACTAAGTCACG |  |
| Hsf1-C-pHYG-AID-F | AGGACCCGACAGAGTACAACGATCACCGCCTGCCCAAACGAGCTAAGAAACGTACGCTGCAGGTCGAC | *HSF1* degron C-term tagging |
| HSF1-C-pHYG-AID-R | ATACTATATTAAATGATTATATACGCTATTTAATGACCTTGCCCTGTGTATCGATGAATTCGAGCTCG |  |
| HSF1 Cterm conf 1 F | TGACCACAGTTATTCCACC | Confirmation of *HSF1* tagging |
| HSF1 Cterm conf 1 R | GCAGTTCAACCTCACTCG |  |
| RPB1-C-pHYG-AID-F | ATTCTCCAAAGCAAGACGAACAAAAGCATAATGAAAATGAAAATTCCAGACGTACGCTGCAGGTCGAC | *RPB1* degron C-term tagging |
| RPB1-C-pHYG-AID-R | AAACTATATATAATGTAATAACGTCAAATACGTAAGGATGATATACTATATCGATGAATTCGAGCTCG |  |
| RPB1 Conf F | ACATCACCTTCTTACTCTCC | Confirmation of *RPB1* tagging |
| RPB1 Conf R | CTTGAAGCTTAGAAGTTGGACG |  |

### Table 4. Primers used for RT-qPCR

| **Name** | **Sequence (5’ 🡪 3’)** |
| --- | --- |
| BTN2 F+598 | GTTTTTGTTATTGGCTGTGGAG |
| BTN2 R+598 | CTTCCTCATGCTTAACTAAACC |
| HSP104 ORF F+1646 | CAG CTG CAA GAT TGA CTG GTA TCC |
| HSP104 ORF R+1799 | CCT GAT CTA GAC AAT CTA ACG GC |
| HSP12 ORF F+100 | ACTGACAAGGCCGACAAGGTCGCTG |
| HSP12 ORF R +211 | CTTGACCTTCAGCGTTATCCTTGC |
| HSP26 ORF F+322 | GTCGTGGTTCCTGGTGTCAAAAGC |
| HSP26 ORF R+432 | ACTCTCTTCATTCAAGGTAGATGG |
| HSP82 UTR F+2134 | AACATCATGGCCTTGAATAGGTTAT |
| HSP82 UTR R+2228 | CATGCAGATGCCCTATTTACATACTT |
| MDJ1 F+799 | CAAACACGCATAAGGTCAGTTGTAG |
| MDJ1 R+799 | GTAATTGTCTTTGCCCTGTTGACC |
| SCR1 F +385 | CGGCCGGGATAGCACATATC |
| SCR1 R +438 | CGCCGAAGCGATCAACTTG |
| SSA4 ORF F+815 | GTC TTC GTC TGC TCA GAC ATC |
| SSA4 ORF R+946 | CCA CTG GCT CCA ATG TAG ATC |
| TMA10 ORF F+ 137 | GCG ATG AGA TTA ATG ACT TAA TCG |
| TMA10 ORF R+ 244 | GCA AAT CAG AAA GCC TTC TTT C |

### Table 5. Primers used for ChIP

| **Name** | **Sequence (5’ 🡪 3’)** |
| --- | --- |
| ACT1 ORF F +811 | GGTTTCTCTCTACCTCACGC |
| ACT1 ORF R +879 | CATCAAGTAGTCAGTCAAATCTCTACC |
| ARS504 F | GTCAGACCTGTTCCTTTAAGAGG |
| ARS504 R | CATACCCTCGGGTCAAACAC |
| HMLα1 ORF F+339 | TGTCTTGTCTTCTCTGCTCG |
| HMLα1 ORF R+473 | GCATAATTATTCGTCAACCACTCTAC |
| HSP12 UAS F -965 | GTCCAGGTGGAGTGCGATTTGTTC |
| HSP12 UAS R -700 | CCTACCTTCTCCCACTTTTCTGTG |
| HSP12 ORF F +100 | ACTGACAAGGCCGACAAGGTCGCTG |
| HSP12 ORF R +211 | CTTGACCTTCAGCGTTATCCTTGC |
| HSP82 UAS F -303 | GTCACATATTGTTCGAACAATTCTGG |
| HSP82 UAS R -238 | CTTCCACGGCGTTCTAGAAAAAAAAG |
| HSP82 UAS R -88 | TGAGGAGGTCACAGATGTTAAGAATT |
| HSP82 ORF F +1248 | GTTCTACTCGGCTTTCTCCAAAAATAT |
| HSP82 ORF R +1444 | CAGCCTTTAGAGATTCACCAGTGATG |
| HSP82 UTR F +2134 | AACATCATGGCCTTGAATAGGTTAT |
| HSP82 UTR R +2228 | CATGCAGATGCCCTATTTACATACTT |
| HSP104 UAS F -266 | CTTAAACGTTCCATAAGGGGC |
| HSP104 UAS R -195 | TGCAGTTCTTTGAGATGGGCC |
| HSP104 ORF F +1646 | CAGCTGCAAGATTGACTGGTATCC |
| HSP104 ORF R +1799 | CCTGATCTAGACAATCTAACGGC |
| PGM2 ORF F +914 | TCGTTTCTCCAGGTGACTCC |
| PGM2 ORF R +1038 | AACACGGTCTATGGCTCCTG |
| SSA4 UAS F -396 | ACACGAAAGATATCTCAACTCTAGCC |
| SSA4 UAS R -145 | TGTTACTGTCGTCAAACTAAGGAG |
| SSA4 ORF F +815 | GTCTTCGTCTGCTCAGACATC |
| SSA4 ORF R +946 | CCACTGGCTCCAATGTAGATC |
| SSA4 UTR F +1762 | GAGGAATACAAGGAAAGGCAAAAG |
| SSA4 UTR R +2079 | TTAAACTCTGGCTTATGACGATGAG |
| TMA10 UAS F -302 | TGGTGGTATACGAGATAGTGATAG |
| TMA10 UAS R -185 | GTGGTGTCTGGATTACCTACCTCAAG |
| TMA10 ORF F +137 | GCGATGAGATTAATGACTTAATCG |
| TMA10 ORF R +244 | GCAAATCAGAAAGCCTTCTTTC |
| TUB1 ORF F+1101 | GTTCAAAGAGCTGTCGAGCAGG |
| TUB1 ORF R+1190 | GTAGCAAATACCGATCTTGAAACC |
| YFR057W F+357 | GCATACATATGAATATTACAACCAC |
| YFR057W R+459 | GCATTATGGCTTTGTTACG |

### Table 6. Primers used for Taq I - 3C

| **Name** | **Sequence (5’ 🡪 3’)** |
| --- | --- |
| ARS504 F | GTCAGACCTGTTCCTTTAAGAGG |
| ARS504 R | CATACCCTCGGGTCAAACAC |
| BTN2 F+1535 | CAAGTGATTCACAAAATACAAGC |
| BTN2 R+1535 | GTTGAATTCTATTCAAGCAGAAATG |
| HSP12 F-47 | ACGTATAAATAGGACGGTGAATTGC |
| HSP12 R-47 | TTCAGAAGCTTTTTCACCGAATC |
| HSP12 F+279 | AAAAGGCAAGGATAACGCTGAAG |
| HSP12 R+279 | AAACCATGTAACTACAAAGAGTTCCG |
| HSP26 F+221 | TATGATCCCAGAGATGAAACC |
| HSP26 R+221 | GAAACCGAAACCAGATGG |
| HSP26 F+807 | GAATTAGTTTAGATATATATGAGTTGATG |
| HSP26 R+807 | ACGAGGTTAGATTCCTTCG |
| HSP82 F+740 | AATTAGTCGTCACCAAGGAAGTTG |
| HSP82 R+740 | AATGCTTAACGTACAATGGGTCTTC |
| HSP82 F+1446 | GCCAGAACACCAAAAGAACATCTAC |
| HSP82 R+1446 | ATTCATCAATTGGGTCGGTCAAG |
| HSP82 F+2189 | ATGAGGATGAAGAAACAGAGACTGC |
| HSP82 R+2189 | ACACACTAGACGCGTCGGAATAG |
| HSP104 F-63 | AGGCATTGTAATCTTGCCTCAATTC |
| HSP104 R-63 | ATCGTTAGAGCCCTTTCTGTAAATTG |
| HSP104 F+782 | GTAAGACCGCTATTATTGAAGGTG |
| HSP104 R+782 | TTCTTCGATTTCCTTCAAAACACC |
| HSP104 F+1550 | CCCTTGATGCTGAACGTAGATATG |
| HSP104 R+1550 | CCACATTTTGGATCATGGAGTTG |
| HSP104 F+1922 | TTTAGCTAATCCAAGGCAACCAG |
| HSP104 R+1922 | ACCTTCATCGTACCCGACATAAC |
| HSP104 F+2756 | AGGTGATGACGATAATGAGGACAG |
| HSP104 R+2756 | TCTTTTGCTCGGGTGTCAAGTTC |
| MDJ1 F+605 | ATTTTGAAGACCTGTTTGGTGCTG |
| MDJ1 R+605 | AGCGCAGAGAATCTTAACTGAACG |
| MDJ1 F+799 | CAAACACGCATAAGGTCAGTTGTAG |
| MDJ1 R+799 | GTAATTGTCTTTGCCCTGTTGACC |
| MDJ1 F+1729 | AGATAAGGAAAAATAGAAGCCTTTACG |
| MDJ1 R+1729 | GTCAAAACCTTTAGCGTCTCAGTG |
| SSA2 F+198 | AGGTAACAGAACCACTCCATCTTTC |
| SSA2 R+198 | GCTTCATATCACCTTGGACTTCTG |
| SSA2 F+1368 | TCTCTACTTATGCTGACAACCAACC |
| SSA2 R+1368 | TTCAATTTGTGGGACACCTCTTG |
| SSA4 F+198 | GCCTTCTTATGTGGCTTTTACTGAC |
| SSA4 R+198 | TTTACGTCCGATCAGACGCTTAG |
| SSA4 F+1079 | TGCTGATTTGTTTAGATCTACATTGG |
| SSA4 R+1079 | TAATACCACCTGCAGTTTCAATACC |
| SSA4 F+2255 | ATAAGAAAGTCATCGCCAAACAAC |
| SSA4 R+2255 | GTGTTAAACTCCGGTCAAAAGAAAC |
| TMA10 F+159 | ACGAAGCAAAGTCTAACCCAAAG |
| TMA10 R+159 | TTTCATTGTTTTGGGAGTTAGAGC |
| TMA10 F+812 | ATGCAAAAACACTTCCCAGAATAG |
| TMA10 R+812 | CCGGTTATAGGACCCTTATTGATG |
